# Human IL-6 fosters long-term engraftment of patient derived disease-driving myeloma cells in immunodeficient mice

**DOI:** 10.1101/2024.01.21.576547

**Authors:** Zainul S. Hasanali, Alfred L. Garfall, Lisa Burzenski, Leonard D. Shultz, Yan Tang, Siddhant Kadu, Neil C. Sheppard, Derek Dopkin, Dan T. Vogl, Adam D. Cohen, Adam J. Waxman, Sandra P. Susanibar-Adaniya, Martin Carroll, Edward A. Stadtmauer, David Allman

## Abstract

Multiple myeloma is a largely incurable and life-threatening malignancy of antibody-secreting plasma cells. An effective and widely available animal model that recapitulates human myeloma and related plasma cell disorders is lacking. We show that busulfan-conditioned hIL-6 transgenic NSG mice (NSG+hIL6) reliably support the engraftment of malignant and pre-malignant human plasma cells including from patients diagnosed with monoclonal gammopathy of undetermined significance, pre- and post-relapse myeloma, plasma cell leukemia, and AL amyloidosis. Consistent with human disease, NSG+hIL6 mice engrafted with patient-derived myeloma cells, developed serum M spikes, and a majority developed anemia, hypercalcemia, and/or bone lesions. Single cell RNA sequencing showed non-malignant and malignant cell engraftment, the latter expressing a wide array of mRNAs associated with myeloma cell survival and proliferation. Myeloma engrafted mice given CAR T-cells targeting plasma cells or bortezomib experienced reduced tumor burden. Our results establish NSG+hIL6 mice as an effective patient derived xenograft model for study and preclinical drug development of multiple myeloma and related plasma cell disorders.

## Introduction

Multiple myeloma (MM) and related clonal bone marrow (BM) plasma cell dyscrasias (PCDs) cause ∼100,000 deaths/year worldwide (1). In addition to MM, these disorders include a pre-malignant state called monoclonal gammopathy of undetermined significance (MGUS) (2), a highly aggressive and therapy resistant leukemia termed plasma cell leukemia (PCL) (3) and AL amyloidosis, which is characterized by the formation of monoclonal antibody-driven amyloid fibrils (4). Despite significant recent advances in therapy options for MM, PCL, and AL amyloidosis patients that build on the previous success of proteasome inhibitors and thalidomide analogs (5, 6), the majority of patients experience relapse and eventually succumb to complications of treatment-refractory disease (7).

A major roadblock to curative drug development for myeloma and other PCDs has been the lack of a flexible and readily accessible animal model that recapitulates human disease. In principle, any such model would support the long-term persistence and growth of primary patient derived PCDs in a manner that mirrors both the growth properties of PCDs and key clinical signs such as anemia, hypercalcemia, renal damage, and bone destruction. Currently, the standard approach to study novel therapeutics *in vivo* is in immunodeficient mice engrafted with MM cell lines (8–10). However, cell line xenograft models fail to reliably recapitulate many aspects of clinical disease, do not faithfully model drug resistance mechanisms, and the cell lines used often grow aggressively, in contrast to most slower growing PCDs (11, 12). An alternative approach involves engraftment of human fetal or rabbit bone chips implanted into immunodeficient mice; however, these systems fail to drive clinical signs of disease, and the bone-resident MM cells do not disseminate throughout the skeleton as the disease does in humans (13).

Two patient derived xenograft models have been reported for primary myeloma. The first uses NSG mice in a similar approach to that presented here (14). However, prolonged engraftment, characterization of engrafted cells, characterization of clinical phenotypes and evaluation of cellular immunotherapies have not been performed. The second, from Das *et al.,* showed that immunodeficient mice (RAG2^-/-^ *γ*c^-/-^) harboring humanized versions of several cytokines including G-CSF, GM-CSF, IL3 and IL6 (MISTRG6) afford robust engraftment of patient PCDs (15). Though Das *et al*. determined that IL-6 is essential for PCD engraftment, the necessity of the other humanized genes was not firmly established. This more complex model is also difficult to obtain due to licensing restrictions and requires continuous antibiotic administration, resulting in limited use within the myeloma research community.

We studied the engraftment and long-term persistence of all major PCDs after transfer into transgenic NSG mice harboring a bacterial artificial chromosome (BAC) containing the human IL-6 gene (NSG+huIL6). We reasoned that increasing systemic IL-6 levels with a humanized BAC might be advantageous, because the BAC is likely to contain cis-regulatory elements needed for proper cell-type restricted IL-6 expression and because mouse IL-6 does not stimulate the human IL-6 receptor (16). Our results establish NSG+hIL6 mice as a straightforward and readily accessible system for the study of a wide range of PCD disease manifestations and therapies including newly diagnosed and relapsed myeloma.

## Results

NSG+hIL6 mice were generated by microinjecting a BAC containing the promoter and gene elements of the human IL6 gene on chromosome 7 into fertilized embryos of NSG mice (17). Because heterozygous females had low fertility, we bred males with normal NSG females; ∼50% of the resulting pups carried the BAC. ELISA analyses showed that the majority of NSG+hIL6 mice possessed human IL-6 (mean 246.3 pg/mL, range 0-1020) in sera. hIL-6 levels distributed into two groups, 9-300pg/mL and 300-600pg/mL (**Figure 1**). There were no associations or trends observed in downstream experiments between the two groups. These IL-6 levels are higher than those observed in normal human sera (<5pg/mL) (18), yet they are in the range of IL-6 expression in PCDs (0.01-4ng/mL) (19).

**Figure 1:**
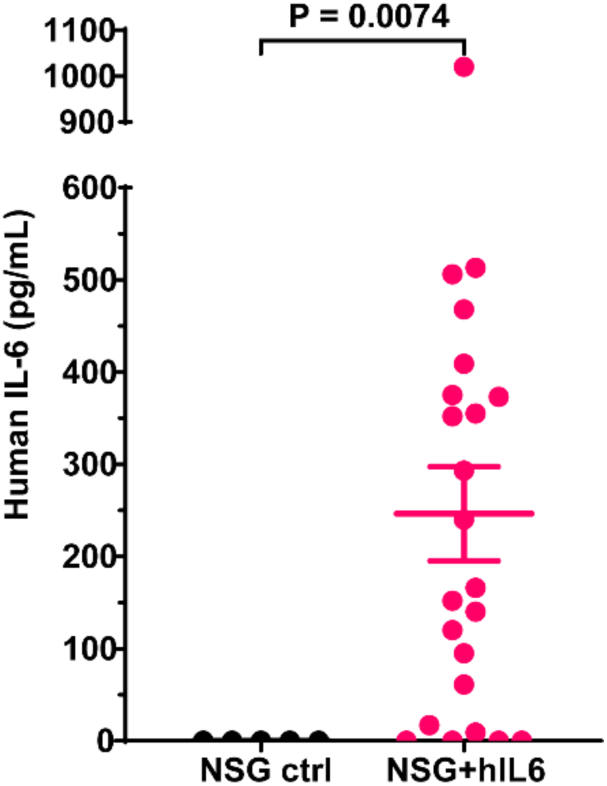
Human IL-6 in NSG+hIL6 sera. Sera from 12–20-week-old NSG (n=5) and NSG+hIL6 (n=23) mice were evaluated for human IL-6 levels by quantitative ELISA. Horizonal lines and error bars indicate the mean and the standard deviation of the mean, respectively.

### Patient MM cell engraftment in NSG+hIL6 mice

Inspired by established xenotransplantation protocols (20), we examined the impact of host pre-conditioning with busulfan with and without the presence of the hIL6 locus on engraftment of primary MM cells following intraosseous injection of patient BM mononuclear cells. Initially we tested for engraftment of malignant plasma cells from two new diagnosis MM patients (MM1 and MM2) following transfer of 1x10^6^ mononuclear BM cells per mouse. We evaluated human antibodies in sera every 5 weeks out to 20 weeks post-transfer and then at 52 weeks post-transfer. Within 5 weeks there was clear detection of human Ig in sera in busulfan treated NSG+hIL6 mice for both MM1 and MM2. By contrast, at this time, engraftment was far less routine for busulfan treated NSG mice and NSG+hIL6 mice without busulfan (**Figure 2A**). Furthermore, for most NSG+hIL6 mice, serum titers for human Ig increased progressively over 20 weeks (**Figure 2B**). Whereas the majority of NSG+hIL6 mice exhibited clear signs of engraftment within 5-10 weeks, by 20 weeks post injection many NSG mice also scored positive for human Ig serum antibodies (**Figure 2C**). Serum protein electrophoresis (SPEP) gels revealed a gamma region M-spike for 9 tested xenografted mice at 15 weeks post injection that was absent in a non-xenografted control (**Figure 2D**). Also consistent with engraftment of monoclonal plasma cells, ELISA for human heavy chains IgG, IgM or IgA showed the presence of only IgG, reflecting the same clonality of the engrafted myeloma clone from the original patient sample (**Figure 2E**). Staining of BM tissue sections with anti-human CD138 and kappa light chain antibodies revealed clusters of human plasma cells (**Figure 2F**). Flow cytometric analyses of BM cells from serum IgG^+^ NSG+hIL6 mice implanted from MM2 showed Igκ restricted light chain expression **(Figure 2G left panel)**, in line with the engrafted myeloma clone. The fraction of all BM cells that were human myeloma cells ranged from <1-12±4%. Consistent with the slow growth rate of malignant plasma cells, under 3% of myeloma cells derived from MM2 engrafted mice were Ki67^+^ (21) (**Figure 2G right panel**). We conclude that NSG+hIL6 mice provide a supportive environment for the efficient engraftment and long-term persistence of primary MM cells.

**Figure 2:**
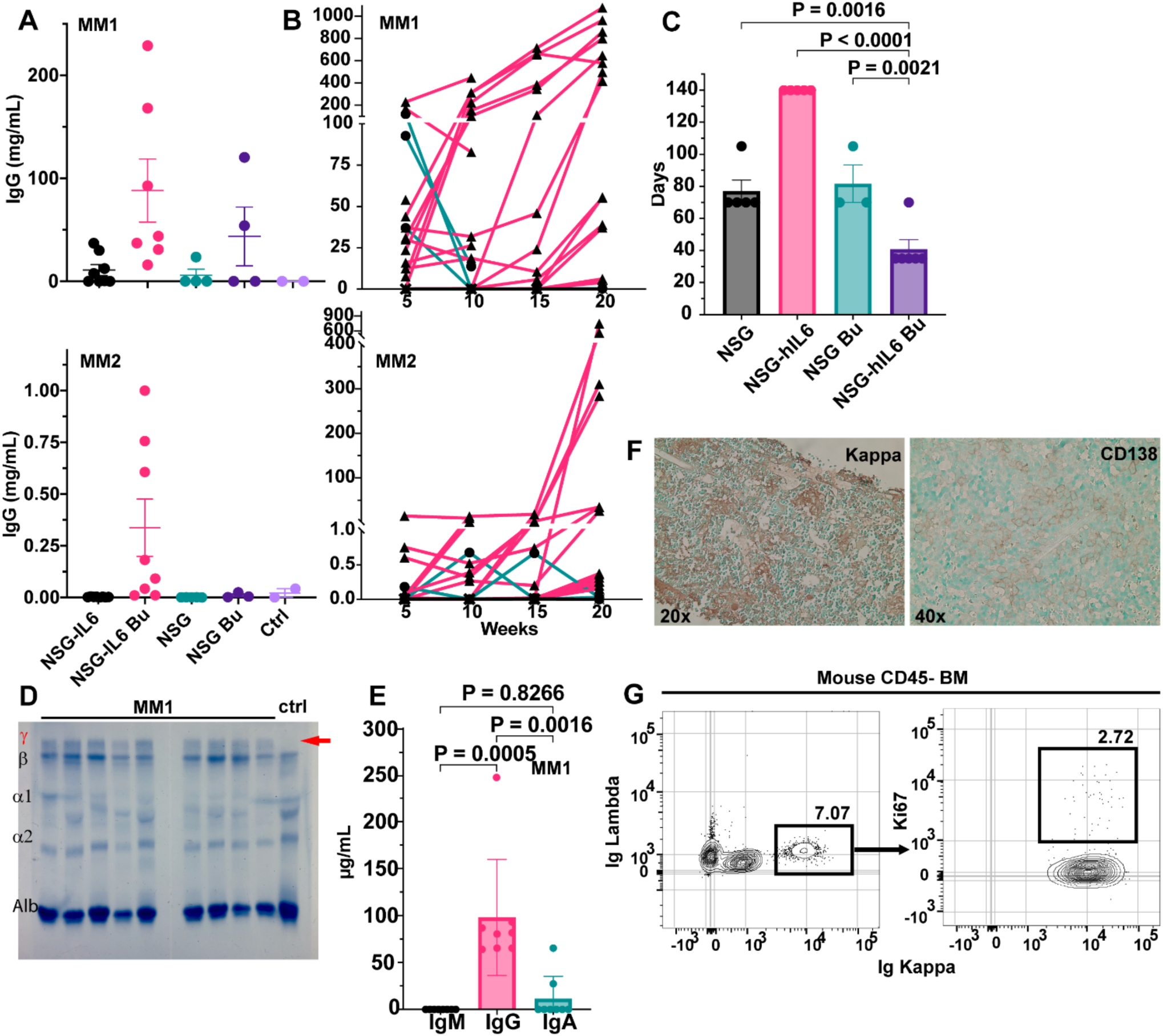
NSG+hIL6 mice support primary patient MM. BM cells from one of two newly diagnosed MM patients (“MM1” and “MM2”) were transferred via intraosseous injection to NSG or NSG+hIL6 adults with and without busulfan pretreatment. (A) Sera from the indicated cohorts were evaluated for human IgG levels by ELISA 5 weeks post injection. “Ctrl” indicates saline injected mice. Horizonal lines and error bars indicate the mean and standard deviation of the mean, respectively. (B) Serum IgG levels in MM1 and MM2 engrafted mice over 20 weeks grouped by engraftment status (unengrafted: green line and circle, engrafted: pink line and triangle). (C) Time to engraftment for NSG vs NSG+hIL6 hosts with or without preconditioning with MM1 (NSG-hIL6 Bu vs NSG (p=0.120), NSG-hIL6 (p=0.082), NSG Bu (p=0.224)). Each data point represents a single mouse. (D) SPEP analysis of 9 sera samples from mice engrafted with MM1 vs an unengrafted control. Gamma region denoted with the red ψ. Red arrow denotes M-spike representative of myeloma engraftment. (E) Total IgM, IgG and IgA serum levels from 7 mice engrafted with MM1 were determined by ELISA. (F) Histologic sections prepared from the BM of an MM1 engrafted NSG+hIL6 host were stained with antibodies specific for human Ig kappa or CD138. (G) BM cells from an MM2 engrafted NSG+hIL6 host were stained with anti-mouse CD45, fixed and permeabilized, then stained with anti-human Ig kappa, Ig lambda, and anti-Ki67 antibodies before flow cytometric analysis. Plots are pre-gated on viable mouse CD45^-^ cells and the right-most plot is gated on the kappa positive myeloma population. For (C) and (E) columns and error bars indicate the mean and standard deviation of the mean, respectively.

### Engraftment of a spectrum of PCDs

Next, we asked whether NSG+hIL6 support engraftment of other PCDs. Using the same methodology, we were able to engraft nearly 100% of NSG+hIL6 mice with samples from donors experiencing MGUS, smoldering MM, *de novo* MM, relapsed/refractory (R/R) MM, PCL and AL amyloidosis (**Figure 3A**). Additionally, we achieved engraftment from three cryopreserved patient samples taken from relapsed MM or PCL patients some 5 years earlier (**Figure 3A, asterisks**), indicating a flexibility in model setup not shared by other systems.

**Figure 3:**
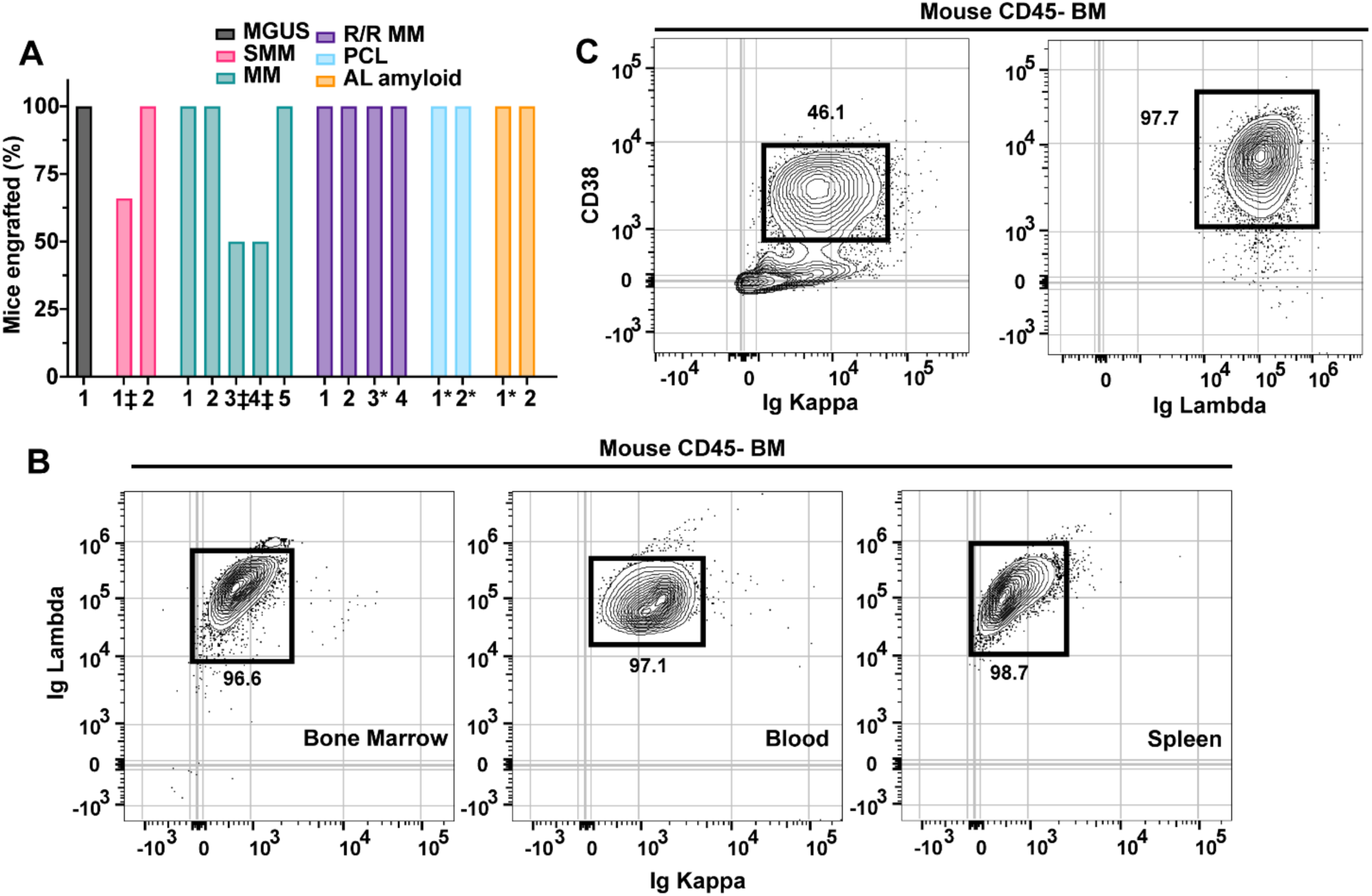
NSG+hIL6 mice support major plasma cell dyscrasias. NSG+hIL6 mice served as hosts for BM cells derived from patients with MGUS, smoldering multiple myeloma (SMM), newly diagnosed multiple myeloma (MM), plasma cell leukemia (PCL), relapsed/refractory myeloma (R/R MM) or AL amyloidosis (AL amyloid). (A) Shown is the fraction of mice in each cohort with sera scoring positive for human IgG patients at 10 weeks post-transfer. (n=5 hosts/grp). *Recipients of previously frozen human BM cells. ‡ Samples not reaching 100% engraftment were prematurely terminated after 3 weeks due to mycoplasma contamination. (B) Flow cytometric analysis for Ig lambda and Ig kappa expression in permeabilized mouse BM (left), blood (middle) and spleen (right) cells harvested from an NSG+hIL6 mouse engrafted with BM from a PCL patient. (C) Analysis of CD38 and Ig kappa or Ig lambda expression for mouse BM cells from separate NSG+hIL6 hosts engrafted previously with BM cells from the MM2 donor (left) or the PCL patient illustrated in (B). For (B) and (C) plots were gated on viable mouse CD45 negative singlets.

Flow cytometric analysis of BM from Igα^+^ PCL engrafted mice showed Igα restricted light chain expression on the BM engrafted clone **(Figure 3B, left panel)**. Additionally, Igα^+^ restricted cells dominated the blood **(Figure 3B, middle panel)** and were noted in spleen **(Figure 3B, right panel)**. Circulating disease was only detectable in mice engrafted with BM cells from a PCL patient, not other PCDs, in line with observed human phenotypes. Also of note, whereas we often detected surface expression of the ectoenzyme and drug target CD38, CD38 levels varied on the plasma cells derived from different donors (**Figure 3C**). We conclude that the BM microenvironment of NSG+hIL6 mice supports the engraftment of a wide variety of PCDs with similar disease-affiliated characteristics.

### Single cell RNAseq (scRNAseq) analyses

Because we engrafted unsorted BM mononuclear cells from patients with PCDs, we sought to further characterize human cells engrafted into NSG+hIL6 hosts. We performed single cell RNAseq (scRNAseq) on total BM cells from an NSG+hIL6 mouse 52 weeks after implantation with mononuclear BM cells from a patient with IgG lambda R/R MM with t(4;14), sample MM3. We utilized the Parse Biosciences pipeline to prepare and analyze data. Human and mouse cells were distinguished by the presence of species-specific mRNA transcripts. As shown in blue and green (**Figure 4**), human cells comprised a small fraction of total BM cells and segregated into three clusters. These cells included a cluster containing clonal human plasma cells denoted by mRNAs for the IGHG1 and IGL2 genes, the myeloma and plasma cell transcription factors BLIMP1 (22) and IRF4 (23), and the myeloma-associated proteins CD38 (24), CD200 (25), FGFR3 (Fibroblast Growth Factor Receptor-3) and NSD2 (Nuclear receptor binding SET Domain protein-2) (26), the latter two resulting from the t(4;14) translocation present in this patient’s myeloma. Additionally, we detected human T cells (CD2^+^ CD3ε^+^) and mast cells (c-Kit^+^ GATA2^+^, IgE Fc receptor subunit β^+^). T cells were enriched for transcripts for immune quiescence (TIGIT, LAG3, PD1), and, notably, no graft vs host disease was observed. No human CD34^+^ stem cell, B-cell (IgM, IgD, PAX5, CD20, CD19), macrophage (CD16, CD14), neutrophil (MPO), megakaryocyte (TPO), stromal cell (FN1, FGFR2), osteoblast (BGLAP, SPP1) or endothelial cell (CDH5, MCAM) specific markers were detected. Altogether, based on the results in Figures 1-4 we conclude that NSG+hIL6 mice support the efficient and long-term engraftment of primary PCDs.

**Figure 4:**
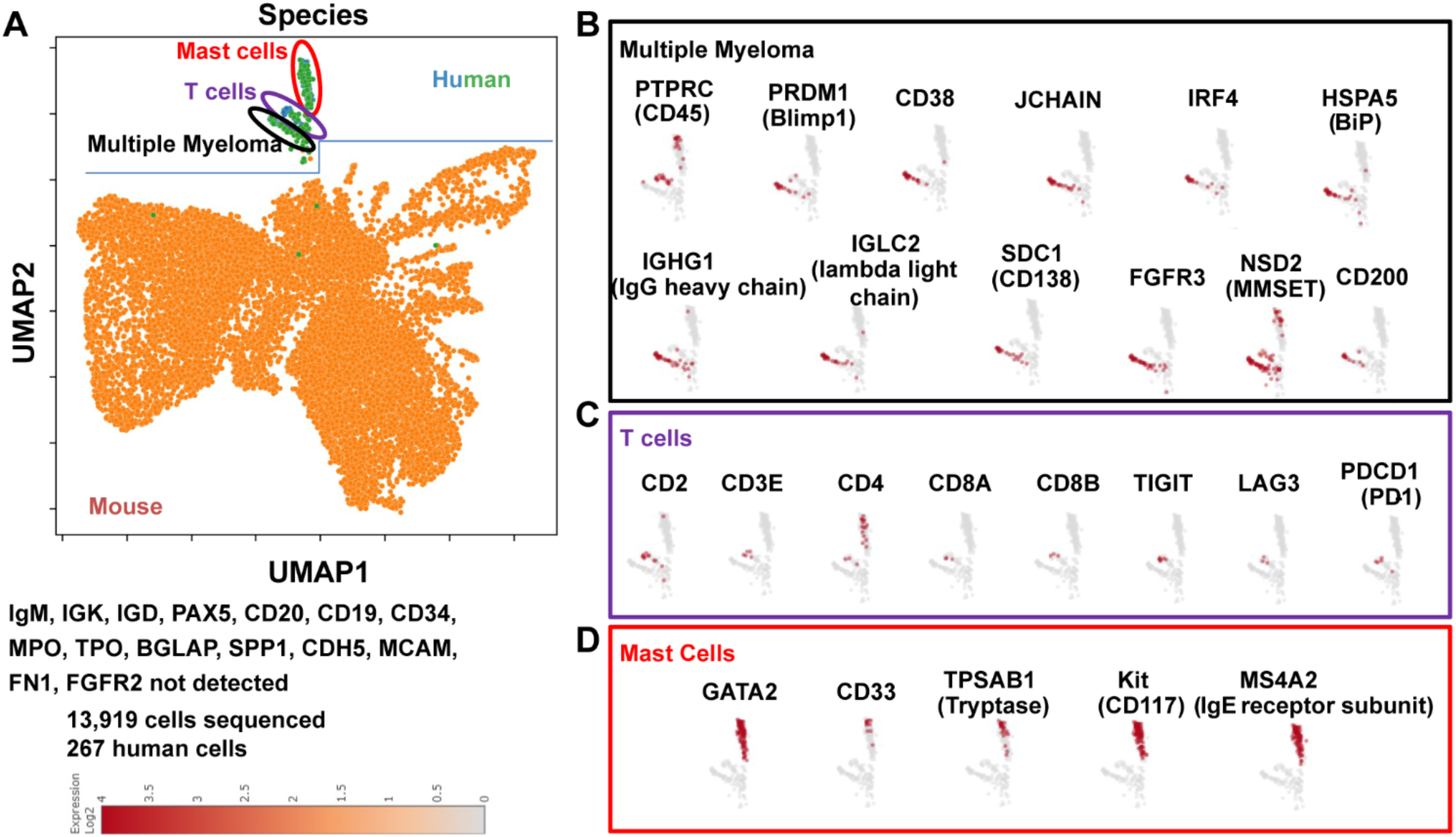
Characterization of NSG+hIL6 mice engrafted mice. BM from an NSG+hIL6 mouse engrafted with mononuclear human BM cells from patient sample R/R MM3 was isolated 52 weeks after intraosseous injection and subjected to scRNAseq using the Parse Biosciences processing and analysis pipeline. (A) UMAP denotes the presence of mouse cells (orange) and human cells (green and blue). Human cells form 3 clusters. Gene expression profiles define these as (B) myeloma cells (black), (C) T cells (purple) and (D) mast cells (red).

### Myeloma engrafted NSG+hIL6 mice exhibit signs of disease

To test the utility of NSG+hIL6 mice for study of MM-associated disease states, we probed for signs of urine Ig, anemia, hypercalcemia, MM cell dissemination throughout the skeleton, and bone destruction in mice engrafted with cells from the MM1 or MM2 donor. Due to logistic reasons, not all mice were able to be tested for all clinical sequelae of disease. At 15 weeks post injection, urine from several engrafted mice possessed detectable titers of human Ig (**Figure 5A**), similar to many patients with MM. Likewise, RBC counts were significantly lower in serum IgG^+^ mice compared to non-engrafted controls (**Figure 5B**). Third, though not common, mice with high ionized serum calcium levels were detected in IgG^+^ mice at levels well above those of non-engrafted mice (**Figure 5C**). Fourth, whereas all mice were inoculated into their left femur, at 8 weeks post-transfer, Igκ^+^ MM cells were readily detected in both the left (**Figure 5D middle**) and the right femur (**Figure 5D right**), confirming spread within the skeleton, a hallmark of MM.

**Figure 5:**
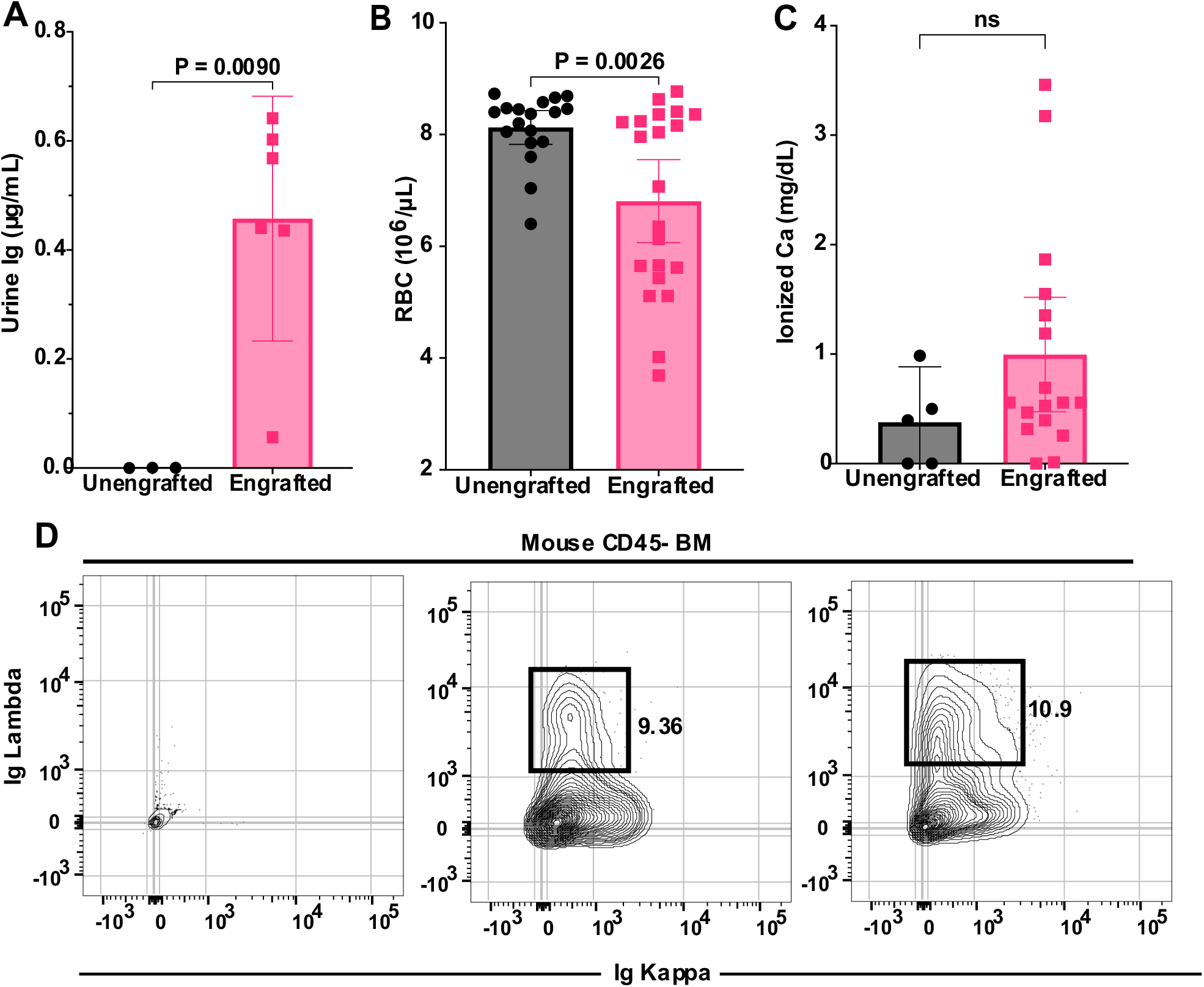
Myeloma engrafted NSG+hIL6 mice with sequelae of human disease. (A) Urine from 6 NSG+hIL6 mice engrafted 15 weeks previously with MM1 BM cells and 3 unengrafted controls was evaluated for human Ig. (B) RBC counts from engrafted vs. unengrafted mice at 15 weeks post injection. (C) Serum ionized calcium concentrations in engrafted (n=15) compared to unengrafted controls (n=5) at 15 weeks. (D) Flow cytometric analysis of Ig kappa and Ig lambda expression for permeabilized BM cells from an unengrafted NSG+hIL6 mouse (left plot),the left femur (middle plot) and right femur (right plot) of a serum human IgG^+^ NSG+hIL6 mouse given MM1 BM cells 12 weeks previously. BM cells were only injected into the left femur. Columns and error bars indicate the mean and standard deviation of the mean, respectively.

At 52 weeks, several engrafted mice were assessed for skeletal abnormalities by microCT scan prior to euthanasia. These mice showed thinned bone with vertebral lesions, sternal lesions and even a fractured femur (**Figure 6**). All of these clinical manifestations are commonly observed in advanced human myeloma (27). Together these data indicate that the NSG+hIL6 xenograft model also recapitulates the clinical sequelae of human MM, a feature that heretofore has not been described in other models. Lastly, majority of engrafted mice succumbed between ∼100 and 400 days post-transfer and eventually all mice died. Except for one mouse, all mice died only after detection of circulating Ig, indicating myeloma was responsible for death. The median overall survival of MM1 and MM2 was 296 and 361 days, respectively (**Figure 7**). When cause of death was analyzed, 11 (24%) mice had hind limb paralysis, 16 (35%) became moribund and 14 (30%) were found dead in their cage (**Table 1**).

**Figure 6:**
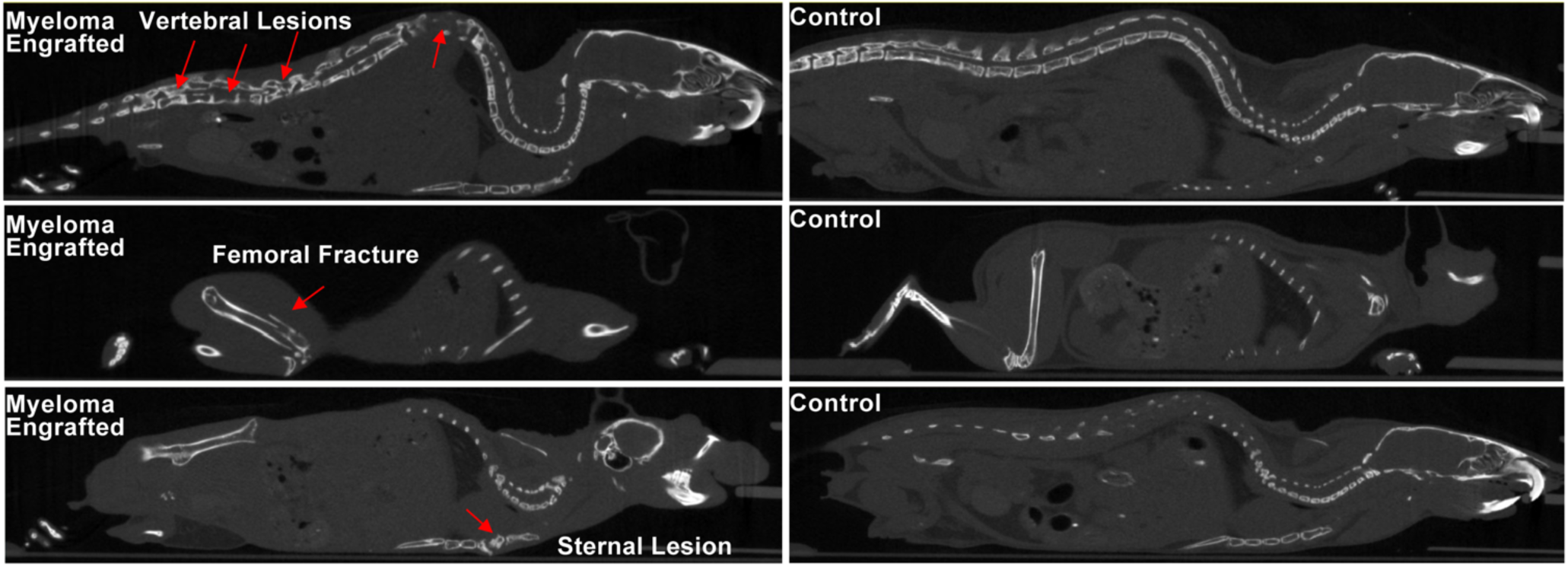
Myeloma engrafted NSG+hIL6 mice develop skeletal lesions. Computed tomography (CT) scans of surviving human IgG^+^ NSG+hIL6 were performed at 52 weeks post injection. Vertebral (top left), femoral (middle left) and sternal (bottom left) lytic lesions in MM1 and MM2 engrafted mice (red arrows) were noted compared to unengrafted mice.

**Figure 7:**
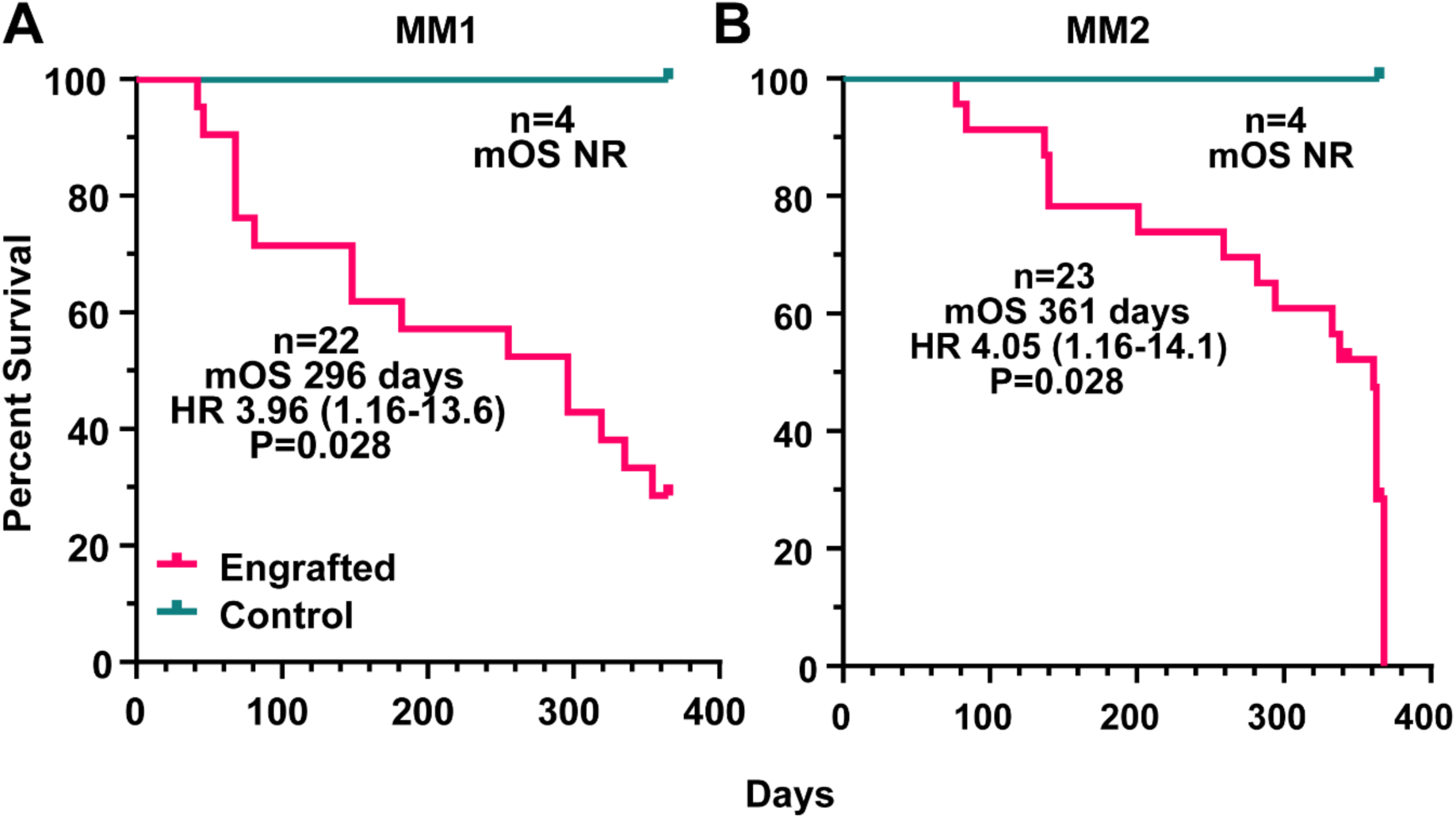
Mortality of myeloma engrafted NSG+hIL6 mice. Kaplan-Meier curves for NSG+hIL6 mice that were engrafted at 16-weeks of age with BM cells from donor MM1 (A) or MM2 (B) or non injected (control). All mice were monitored for humane endpoints over the indicated time frames. There was only a single mouse, within the MM1 cohort, that was injected with myeloma cells and died before IgG was detectable in the serum. All others had detectable IgG at the time of death.

**Table 1:** Characteristics of MM1 and MM2 engrafted NSG+hIL6 mice.

| MM1 |  |  |  |  |  |  |  |
| --- | --- | --- | --- | --- | --- | --- | --- |
|  | Cause of death | Survival (days) | Sex | Hypercalcemia | Urine light chains | Anemia | Bone lesions (~52 weeks) |
| NSG |  |  |  |  |  |  |  |
| 2428 | end of study | 397 | F |  |  | no |  |
| 2436 | found dead | 255 | F |  | yes | no |  |
| 2439 | moribund | 255 | M |  | no | no | yes |
| 2444 | hind limb paralysis | 255 | F | no |  | no |  |
| NSG IL6 |  |  |  |  |  |  |  |
| 2412 | moribund | 148 | M |  |  | no |  |
| 2414 | censored, death with anesthesia |  | M |  | yes | no |  |
| 2417 | moribund | 255 | M |  |  | yes |  |
| 2418 | found dead | 319 | F | no |  | no |  |
| 2421 | moribund | 296 | F |  |  | no |  |
| 2437 | moribund | 182 | F | no |  | yes |  |
| 2443 | found dead | 81 | F |  |  |  |  |
| 2447 | found dead | 354 | F | yes |  | no |  |
| NSG Busulfan |  |  |  |  |  |  |  |
| 2416 | moribund | 51 | M |  |  |  |  |
| 2420 | hind limb paralysis | 296 | F |  |  | no |  |
| 2435 | hind limb paralysis | 335 | F |  |  | yes | yes |
| 2442 | found dead | 46 | F |  |  |  |  |
| NSG IL6 Busulfan |  |  |  |  |  |  |  |
| 2411 | found dead | 42 | M |  |  |  |  |
| 2413 | moribund | 148 | M |  |  | yes |  |
| 2415 | moribund, massive ascites | 68 | M |  | no |  |  |
| 2427 | hind limb paralysis | 397 | F |  | yes | yes |  |
| 2438 | moribund | 68 | F | no |  |  |  |
| 2441 | moribund | 296 | M |  | yes | yes |  |
| 2445 | moribund | 68 | F |  |  |  |  |

Table 1: Characteristics of MM1 and MM2 engrafted NSG+hIL6 mice (continued)
| MM2 |  |  |  |  |  |  |  |
| --- | --- | --- | --- | --- | --- | --- | --- |
|  | Cause of death | Survival (days) | Sex | Hypercalcemia | Urine light chains | Anemia | Bone lesions (~52 weeks) |
| NSG |  |  |  |  |  |  |  |
| 2802 | hind limb paralysis | 137 | M |  | yes | no |  |
| 2804 | hind limb paralysis | 201 | M |  |  | yes |  |
| 2810 | end of study | 397 | F |  |  | no |  |
| 2813 | moribund | 140 | M |  |  | no |  |
| 2818 | moribund, massive ascites | 333 | F | yes |  | no | yes |
| NSG IL6 |  |  |  |  |  |  |  |
| 2823 | found dead | 361 | F |  |  | no |  |
| 2832 | found dead | 338 | F |  |  | no |  |
| 2828 | moribund | 363 | M | yes |  | no | yes |
| 2829 | censored, death from anesthesia |  | M |  |  | no |  |
| 2837 | end of study | 397 | M | yes | no | no |  |
| 2825 | found dead | 368 | M | yes |  | no |  |
| 2817 | end of study | 397 | F |  |  | no |  |
| 2821 | found dead | 282 | F | no |  | no |  |
| NSG Busulfan |  |  |  |  |  |  |  |
| 2807 | hind limb paralysis | 294 | F | yes |  | yes |  |
| 2808 | moribund | 77 | F |  |  |  |  |
| 2801 | hind limb paralysis | 333 | M |  |  | no |  |
| NSG-IL6 Busulfan |  |  |  |  |  |  |  |
| 2803 | moribund | 363 | M |  |  | no | yes |
| 2805 | moribund | 84 | M |  | yes |  |  |
| 2806 | hind limb paralysis | 363 | F |  |  | yes | yes |
| 2809 | hind limb paralysis | 363 | F |  |  | no | yes |
| 2811 | end of study | 397 | F |  | yes | yes |  |
| 2812 | moribund | 140 | F |  |  | yes |  |
| 2814 | hind limb paralysis | 259 | M |  |  | yes |  |

### Responses to anti-myeloma therapies

To test the usefulness of NSG+hIL6 mice in modeling multiple myeloma therapy, we treated 30-week myeloma-engrafted mice from cohort MM2 with the proteasome inhibitor bortezomib or human BCMA targeted CAR T cells, both of which are MM therapies with proven clinical efficacy. For the latter experiment we employed untransduced (UTD) T-cells as a negative control. For CAR T cells, mice were inoculated with 3x10^5^ cells per dose at 0- and 5-weeks post engraftment, and serum human IgG titers traced weekly over three months. As shown, BCMA directed CAR T-cells prevented the steady increase in IgG level, indicating disease control as compared to control T cells (**Figure 8A**). Additionally, two separate groups of myeloma engrafted mice were treated with saline or bortezomib at 1mg/kg weekly for 4 weeks, a dose and schedule roughly equivalent to that of one cycle of therapy used for human myeloma patients. Serum IgG titers were followed weekly for 6 weeks. Bortezomib significantly decreased titers of human IgG compared to saline controls (**Figure 8B**), indicating the anticipated treatment response. We conclude that NSG+hIL6 mice are a highly suitable option for xenotransplant and study of new drug candidates in human malignant and premalignant plasma cells.

**Figure 8:**
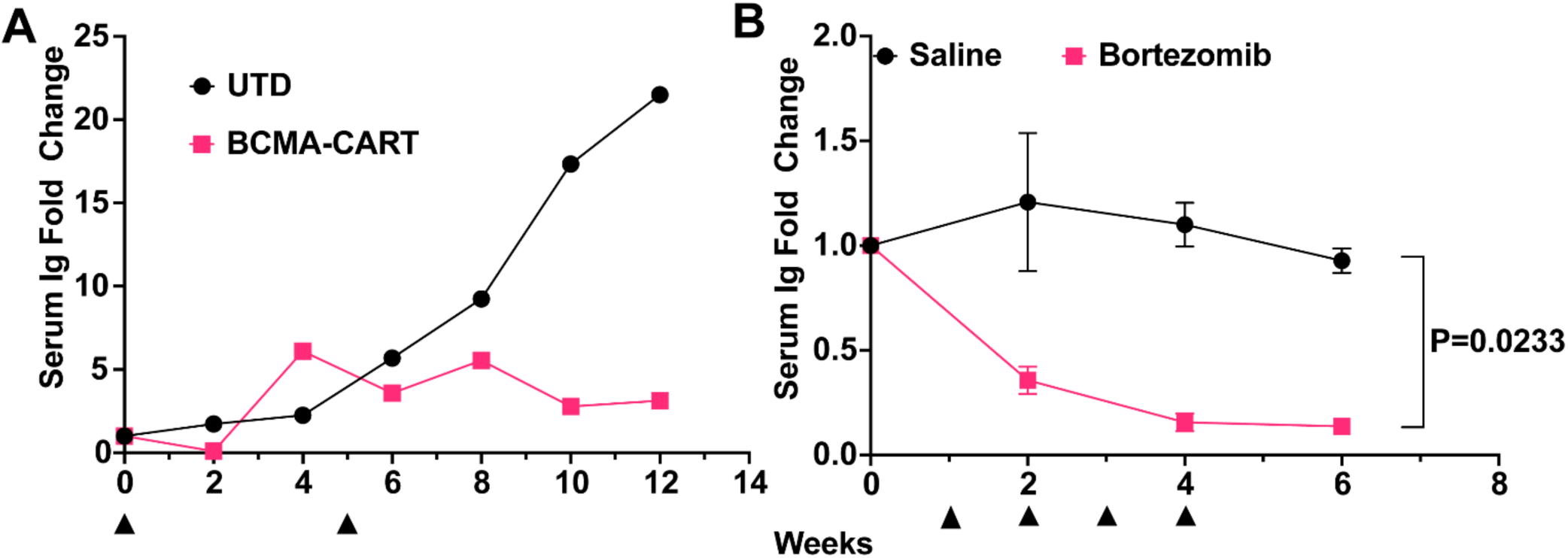
Treatment responses to BCMA directed CAR T cells or bortezomib. (A) NSG+hIL6 mice were implanted with MM1 BM cells. 30 weeks later (“time zero”) serum human IgG positive mice were given 2 doses (black arrowheads) of BCMA CAR T-cells (pink, n=1) or untransduced T-cells (UTD) (black, n=1) (left panel) over a 5-week interval. (B) Separate experiment wherein serum human IgG positive NSG+hIL6 mice were given 4 doses (black arrowheads) of saline (black, n=3, 10uL/g) or bortezomib (red, n=3, 1mg/kg IV) over four weeks.

## Discussion

Our results establish that NSG+hIL6 mice are highly suited for the routine and reproducible engraftment, persistence, and progressive growth of patient-derived malignant plasma cells. Supporting this conclusion, NSG+hIL6 mice were readily engrafted with Ig light chain restricted plasma cells from newly diagnosed and post-relapse myeloma patients as well as donors experiencing MGUS or diagnosed with other plasma cell-driven afflictions including PCL and AL amyloidosis. Further, with time, mouse recipients of myeloma cells experienced progressive increases of human IgG in serum, and many experienced elements of advanced MM such as anemia, hypercalcinemia, bone lesions, and hind limb paralysis consistent with vertebral involvement and cachexia.

Past work has shown that pre-established myeloma cell lines grow rapidly after transfer into NSG mice, often resulting in rapid dominance of host BM within 4 weeks and death soon thereafter (11). By contrast, in NSG+hIL6 mice, patient-derived myeloma cells often comprised a relatively small fraction of all BM cells and appeared to expand relatively slowly, with a median overall survival of 42 or more weeks. Consistent with this conclusion, only small frequencies of Ki67^+^ cells were observed among implanted myeloma cells. The relatively slow growth rates of engrafted plasma cells and the extended survival times of NSG+hIL6 mice are consistent with human disease (21). Indeed, previous attempts to quantify cell division rates for patient myeloma cells suggest relatively slow doubling times ranging from weeks to several months (28, 29). Given that unsorted patient BM mononuclear cells were used for engraftment, we speculate these results suggest that supporting cells may be required for PCD growth in human BM and are either slow growing or altogether absent in many engrafted NSG+hIL6 mice. This hypothesis is further supported by the apparent lack of complete marrow replacement in NSG+hIL6 hosts. scRNAseq of the BM confirmed the presence of the original patient myeloma clone as well as the presence of T cells and mast cells. No other human cell types were detectable by transcripts. Given the presence of mast cells almost a year after myeloma cell engraftment, there are likely CD34+ stem cells that also engrafted, but they were not readily detectable possibly due to sampling error. Future serial transplantation studies may resolve these issues.

Additional facets of the NSG+hIL6 system are also consistent with human MM. In this regard, we note that disparate clinical phenotypes often developed among cohorts of NSG+hIL6 hosts despite receiving identical doses of donor BM cells on the same day from the same myeloma patient. Indeed, some animals took upwards of 6 months before showing detectable antibody in the blood and became moribund soon thereafter, whereas others harbored readily detectable human IgG titers for months before experiencing clinical symptoms. We suggest that these features of the NSG+hIL6 model are relevant because human myeloma phenotypes are similarly variable. In this regard, it remains unknown why certain patients develop certain elements of the disease or why some patients’ disease remains stable for many years before relapsing while others rapidly progress. Ultimately, our model may provide insights into this problem, thereby leading to a better understanding of how myeloma causes complex clinical phenotypes.

With NSG+hIL6 mice we were able to engraft a diverse set of PCDs in >70% of animals from both fresh and frozen samples at 5-10 weeks post injection as compared to NSG mice lacking the human IL6 locus. We used death as a read out, which has seldom been done with past myeloma models, and note that many mice also developed hind limb paralysis at high rates consistent with vertebral involvement and cachexia. Further, longitudinal assay of blood for human antibody titers proved a feasible approach for inferring ongoing treatment response to bortezomib and BCMA CAR-T cell treatment. Further delineation of what cells are responsible for what clinical effects for these and other drugs could lead to development of supportive therapies that prevent myeloma complications in the future.

In summary, we present a new PDX model for PCDs characterized by fidelity to human disease and ease of use. In line with the findings with the MISTRG6 mouse (15), we note dissemination of tumor, circulating disease only with hosts given PCL, and a supportive environment for PCDs in general. The addition of the NSG+hIL6 model and its availability within the research toolbox will aid investigators in the wider PCD research community in the quest for truly durable, curative therapies.

## Methods

### NSG+hIL6 mice

A BAC clone (RP11-469J8) carrying a piece of chromosome 7 with the hIL-6 gene and associated promoter and enhancer elements was microinjected into fertilized NSG embryos. All subsequent breeding involved heterozygous males and wild type females because female NSG+hIL6 mice have low fertility. All mice were bred and maintained under strict clean conditions to minimize risk of infection per protocols within the Penn Stem Cell and Xenograft Core Facility. PCR genotyping for the hIL6 BAC was performed by Transnetyx using the following oligonucleotides: F- GGGAGAGCCAGAACACAGA; R-TGCAGCTTAGGTCGTCATTG.

### Study Approval

All human samples were collected after obtaining informed consent per approved IRB protocol # 842940 through the PCD group at the Hospital of the University of Pennsylvania. All mice experiments were performed under the stem cell and xenograft core IACUC protocol for animal model development. Humane endpoints were used to determine when mice were euthanized. These included weight loss >20%, hind limb paralysis, extreme lethargy and respiratory distress.

### Preparation of primary human cells

All reagents were dedicated to PC isolation to minimize contamination risk. All parts of this procedure except spinning were done in a tissue culture hood with sufficient laminar air flow. 2-5mL of BM aspirate was obtained in green top heparin tubes (not EDTA). Aspirate was diluted to 16mL in DPBS with calcium and magnesium (Thermo) in a sterile 50mL conical tube. 4mL of Ficoll Paque plus (Sigma) was carefully added to the bottom of two 15mL conical tubes, then diluted aspirate was carefully layered over the Ficoll. After equally distributing 8mL of diluted aspirate atop each 4mL Ficoll cushion, tubes were carefully capped and moved to a room temperature swinging bucket centrifuge and spun at 700 RCF for 20 minutes without braking. Buffy coats from both tubes were combined into one 50mL conical. 10mL of DPBS with calcium was added and then mixed with inversion before spinning down at 400 RCF for 5 minutes with normal braking parameters. Supernatant was removed and 5mL of ACK lysis buffer (Thermo) was added. Tube was pipetted up and down and allowed to lyse at room temperature for 5 minutes. Cells were spun down and supernatant removed. Cells were resuspended in 1mL of DPBS, mixed with gentle pipetting until single cell suspension and then counted. For transplantation on the same day, Primocin (Invivogen) was added to the 1mL cell suspension at 100μg/mL along with OKT3 antibody at 10μL/ one million cells (30). Antibiotics and OKT3 treatment were performed to decrease risk of infection from donor pathogens into immunodeficient animals and to deplete GvHD causing T cells, respectively. Cell mixture was resuspended and incubated at 37°C for one hour. 100μL of cell suspension was removed to a 1.2mL eppendorf tube and placed at -20°C for subsequent pathogen testing (IDEXX -hIMPACT panel). Remaining cells were spun down and supernatant removed. Cells were diluted to 1x10^6^ cells/10μL/mouse with an extra 10μL overall to account for loss. Cells were transplanted within 4 hours of cell prep completion.

### Xenograft transplantation

Mice were conditioned with one intraperitoneal injection of busulfan (30mg/kg) 24 hours prior to introduction of prepared patient BM aspirate. Intraosseous injection of aspirate began with anesthetizing mice using isoflurane on anesthesia nose cone. The injection site used was always the left hind limb. The site was shaved just prior to injection and wiped clean using chlorhexidine wipes x3. Meloxicam or Meloxicam SR was injected prior to incision. The mouse’s leg was stabilized in a bent position to allow access to the patellar surface of the femur. A hole is punched through the patellar surface into the shaft of the bone using a 25-gauge needle and then a 30-gauge needle is inserted into the femur. An infusion of 10μL of cells (1x10^6^ cells/mouse) was administered using a small volume syringe. A drop of vet bond was placed at the insertion site when the needle was withdrawn from the femur. Animals were monitored daily for weight loss, malaise, tumors and limb paralysis.

### Following engraftment markers

Blood was the easiest and most reproducible way to follow engraftment of malignant PCs. The NSG mouse has no antibodies at baseline, mouse or human. By following the increase in human titers of total immunoglobulin (Ig) by ELISA it was possible to determine which animals had been engrafted and which had not by ∼5 weeks. In high burden states such as PCL, anticoagulated blood is stainable for malignant cells as well. Blood was collected in Eppendorf tubes and allowed to clot for 30 minutes prior to spinning at 8000 RCF for 8 minutes. Serum was then removed to a new tube leaving red cells behind. Sera was then applied to blood and urine ELISA and SPEP as described.

### ELISA

ELISA plates (Fisher) were coated using 100μL/well coating buffer (NaHCO_3_ 2.93g/L, Na_2_CO_3_ 1.59g/L pH 9.6) and 1μg/mL of unlabeled total anti-human total Ig (Southern Biotech) overnight at 4C or at 37C for one hour. Wells were then washed with wash buffer 3 times (1xPBS with 0.1% Tween 20). Blocking buffer (0.22μM filtered 2% BSA in 1xPBS) was added at 100μL/well and allowed to block at room temperature for 1 hour. 1μL of serum from each mouse was added to a single well at the top of a column. Samples were then serially diluted 1:10 down the columns 3 times for a total of 4 wells per samples. This allowed for 24 samples to be run on one plate. Sera were incubated for one hour. Wells were again washed 3 times with wash buffer. Capture buffer (blocking buffer with 1μg/mL of biotin labelled anti-human total Ig) was added to each well at 100μL/well. Plate was incubated at room temperature for 1 hour and then washed again 3 times. 1μL/10mL of streptavidin-HRP was added to each well at 100μL/well and incubated in the dark at room temperature for 1 hour. Wells were again washed 3 times and plate blotted forcefully against paper towels to remove as much wash buffer as possible. Room temperature TMB substrate (Thermo) was prepared and 100uL added to each well. After wells started turning yellow (1-2 minutes or less), reaction was quenched with 200uL of 1M phosphoric acid. Plates were then assessed for absorbance on a Spectramax microplate reader at 450nm with background subtraction at 570nm. For quantification, Ig kappa or lambda monoclonal protein (Thermo) was run at known concentrations at 10-fold dilutions starting at 1000ng down an entire column (7 dilutions x2 columns). Antibody concentrations were determined using 4PL regression in GraphPad Prism 9.

Urine testing for the presence of Ig was also conducted by ELISA with the same method outlined above. Urine was loaded at 10μL into 100μL wells before dilution due to lower concentration of Ig. ELISA testing for hIL-6 was carried out with an hIL-6 kit (R&D DY206).

### Histology/Immunohistochemistry (IHC)

Tissues were isolated post euthanasia and placed in 10% formalin overnight at 4C. The next day, fixed tissues were removed to two cassettes per mouse, one for soft tissues and one for bones. These cassettes were then placed in 70% Ethanol/30% water and allowed to soak prior to processing. We utilized the histology services of the UPenn veterinary school for standard practices in decalcification, paraffin block embedding, tissue slice preparation and H&E staining. Slices were put on ProbeOn© (Fisher) slides for IHC.

IHC was performed as per IHC protocol (Abcam). After deparaffinization, slides were submitted to sodium citrate buffer antigen retrieval for 30 minutes prior to overnight incubation of primary antibodies (see **Supplemental Table 1**). 10 minutes of 3% hydrogen peroxide was used to reduce endogenous peroxide background before incubation of secondary HRP conjugated antibody and subsequent DAB substrate application for 12 minutes.

### scRNA sequencing of myeloma engrafted BM

BM cells from an NSG+hIL6 mouse engrafted with human myeloma were fixed and stored at -80C with the Evercode cell fixation kit v2 from Parse Biosciences. Just prior to processing, cells were thawed and prepared using the Evercode WT mini v2 kit and associated protocol. This is a plate based barcoding methodology to perform single cell RNA sequencing. Two sublibraries were generated, one with 5000 cells and the second with 10000 cells. Sublibraries were submitted to Azenta Life Sciences for sequencing at equimolar ratios on an Illumina NovaSeq 6000 with paired end 150bp reads (∼350 x 10^6^ reads). Analysis was performed using the Parse Biosciences platform based in R/Python.

### Assaying Serum Ionized Calcium Level

Mouse blood was collected in polypropylene 1.2mL centrifuge tubes without anticoagulants and allow to clot for 30 minutes prior to spinning at 8000 RCF for 8 minutes and removing the sera to a new tube. Sera were then tested for ionized calcium concentration using the Calcium Assay Kit (Abcam 102505).

### Serum Protein Electrophoresis (SPEP)

SPEP was carried out using the QuickGel station from Helena Laboratories and the Split-Beta SPE Kit (3550T) per manufacturer’s instructions.

### Complete Blood Counts

At 15 weeks post injection of patient samples from MM1 and MM2, blood was collected in EDTA coated vacutainer tubes and sent to IDEXX analytics for formal complete blood count testing (test code 375).

### MicroCT scanning

With the help of the Small Animal Imaging Facility Core Resource at UPenn, mice were anesthetized using inductive isoflurane and then maintained through nose cone prior to mounting on the MILabs U-CT ultra-high resolution (∼20 µm) small-to-medium sized animal CT scanner. 4-minute scans were obtained prior to euthanasia. Images were analyzed using ImageJ.

### Flow Cytometry

Cells were isolated from femurs and spleens on ice, lysed for red blood cells using ACK lysis buffer (Thermo) for 5 minutes at room temperature and then stained for live cells with zombie aqua live/dead (Thermo) (10 minutes) and fluorescently labeled antibodies of markers of interest (30 minutes) in 0.1%BSA PBS buffer. Please see **Supplemental Table 1** for antibodies used.

### Statistics

2-sided ANOVA and t-tests were used for comparison or >2 groups or 2 groups respectively and calculated with GraphPad Prism. Significance cut offs were α=0.05. All error bars represent mean ± SD. Cohorts MM1 and MM2 were powered at 80% under the assumption that NSG+hIL6 mice would have an engraftment incidence of 80% based upon observed engraftment in the MISTRG6 model vs. 20% in NSG mice from our prior experience. Treatment with BCMA-CAR T cells and bortezomib could not be powered to the same level given the lack of available myeloma engrafted mice by the time the experiment was begun. Significance was calculated as divergence of slope of least square’s fit of fold change in serum Ig. These latter treatment experiments serve as a proof of concept for study of treatment response in the NSG+hIL6 rather than definitive statements about efficacy of any one specific therapeutic.

### BCMA CAR T-cell generation

CAR T-cells specific for BCMA were kindly provided by the Posey & Milone labs at the University of Pennsylvania. CAR constructs were amplified by PCR, and subcloned into the pTRPE vector before packaging into lentivirus using a VSVG envelope and HEK293T cells. Patient T-cells cells were stimulated and treated with CAR containing lentivirus, then expanded and harvested for injection. BCMA CAR T-cells were given at 3 x 10^6^ cells/mouse.

### Data Availability

Single cell RNA sequencing data has been uploaded to the NCBI GEO database with accession numbers GSE246140, GSM7857100 and GSM785710. Both raw data files as well as normalized files from which the analyses within this manuscript were derived are available for download.

## Author Contributions

ZSH designed and performed experiments, analyzed data, and wrote the manuscript. AG provided patient samples and wrote the manuscript. LB and LDS designed and performed experiments. YT, SK and NCS designed and performed experiments. DD performed experiments. DTV, ADC, AJW and SPS provided patients samples and reviewed the manuscript. MC and EAS designed experiments and wrote the manuscript. DA designed experiments, analyzed data and wrote the manuscript.

## Acknowledgements

We thank members of the Stem Cell and Xenograft Core of the Perelman School of Medicine (SCR_010035), Flow Cytometry and Cell Sorting Core Resource Facility, Human Immunology Core, Small Animal Imaging Facility Core and Comparative Pathology Core and PennVet Comparative Histology Core. This work was supported by a pilot grant from the Abramson Cancer Center Core (P30) (ZSH, DA), the Kohler Drennan Fund for Multiple Myeloma Research (ZH,AG,ES), an ASPIRE award from the Mark Foundation for Cancer Research (DA), the Hematologic Malignancies Translational Center of Excellence (MC, ES), an LLS Scholar award in Clinical Research (AG), National Institutes of Health (NIH) training grant T32-HL07439 (ZH), Amyloidosis Foundation Research Grant (ZH) and NIH grants R01-AI139123 and R21-AI161931 (DA) and CA034196 (LDS). Dr. Carroll receives salary support from VA Merit Award, 5 I01 BX004662.

