## Supplemental Table 1 for "Human IL-6 fosters long-term engraftment of patient derived disease-driving myeloma cells in immunodeficient mice"

Supplemental Information:

Supplemental Table1: Reagent Specifics

|  | **Vendor** | **Product ID** | **Assay** |
| --- | --- | --- | --- |
| Primers |  |  |  |
| F-GGGAGAGCCAGAACACAGA | transnetyx | hIL-6 | genotyping |
| R-TGCAGCTTAGGTCGTCATTG | transnetyx | hIL-6 | genotyping |
| Antibodies |  |  |  |
| OKT3 | Thermo | 16-0037-81 | T-cell depletion |
| anti human total Ig - unlabelebd | Southern Biotech | 2010-01 | ELISA |
| anti human total Ig-Biotin | Southern Biotech | 2010-08 | ELISA |
| anti human IgG - Biotin | Southern Biotech | 2040-08 | ELISA |
| anti human IgM - Biotin | Southern Biotech | 9020-08 | ELISA |
| anti human IgA - Biotin | Southern Biotech | 2050-08 | ELISA |
| sterptavidin-HRP | Biolegend | 405210 | ELISA |
| mouse anti human CD138 | BioRad | MCA2459T | IHC |
| mouse anti human kappa | BioRad | 5268-6010 | IHC |
| anti mouse Ig HRP | Biolegend | 405306 | IHC |
| anti mouse CD45.1 | Biolegend | 110703 | Flow cytometry |
| Streptavidin BUV 661 | BD biosciences | 612979 | Flow cytometry |
| anti human CD138 BUV737 | BD biosciences | 612834 | Flow cytometry |
| anti human  CD19 BUV805 | BD biosciences | 742007 | Flow cytometry |
| anti human  CD38 BV421 | Biolegend | 356617 | Flow cytometry |
| LiveDead Aqua | Thermo | L34957 | Flow cytometry |
| anti human CD47 BV605 | Biolegend | 323119 | Flow cytometry |
| anti human CD20 BV650 | Biolegend | 302335 | Flow cytometry |
| anti human Ki67 BV711 | Biolegend | 350515 | Flow cytometry |
| anti human CD27 BV785 | Biolegend | 302831 | Flow cytometry |
| anti human IgL AF488 | Southern Biotech | 9180-30 | Flow cytometry |
| anti human CD45 PerCP-Cy5.5 | Biolegend | 368503 | Flow cytometry |
| anti human CD269 (BCMA) PE | Biolegend | 357503 | Flow cytometry |
| anti human CD24 PE-CF594 | BD biosciences | 562405 | Flow cytometry |
| anti human CD200 PE-Cy7 | Biolegend | 329211 | Flow cytometry |
| anti human IgK APC | Southern Biotech | 9230-11 | Flow cytometry |
| anti human CD56 AF700 | Biolegend | 362521 | Flow cytometry |
| anti human CD14 APC-Cy7 | Biolegend | 367107 | Flow cytometry |
